# Biogeography and taxonomy of the *Hemileuca maia-nevadensis* Stretch species complex (Lepidoptera: Saturniidae): insights to the midcontinental populations

**DOI:** 10.1101/2022.04.07.487535

**Authors:** Christian Schmidt

## Abstract

An integrated assessment of geographic distribution, genetic variation and interpopulation reproductive barriers of the Great Plains and Great Lakes populations of the *Hemileuca maia-nevadensis* species complex is presented. The taxonomic circumscription of *Hemileuca nevadensis* Stretch, key to understanding the complex evolutionary history of the group, has previously been misinterpreted. Detailed distribution maps based on historic, literature, and contemporary occurrence records, combined with analysis of genetic variation using mtDNA-COI barcode sequences and pheromone compatibility trials, indicate that Great Plains *H*. “*nevadensis*” are disjunct and genetically divergent from nominate *H. nevadensis*, instead being more closely related to Great Lakes populations. Despite the genomic differentiation of the bog buckmoth established in recent studies, similar or identical mtDNA-COI haplotypes indicate past genetic linkages among now-isolated populations across the northern range periphery of the *maia*-complex. Based on the current evidence, northern Great Plains *Hemileuca latifascia* Barnes & McDunnough **stat. nov**. and eastern Great Lakes (bog buckmoth) *Hemileuca menyanthevora* Pavulaan **stat. nov**. (= *Hemileuca iroquois* Dirig & Cryan **syn. nov**.) should be considered as species distinct from others in the *maia*-group. Post-glacial dispersal hypotheses are evaluated in light of molecular variation and ecology, and key knowledge gaps are identified to guide future research.

## Introduction

Members of the *Hemileuca maia-nevadensis* complex have a wide but highly localized distribution across the USA and parts of southern Canada. Despite the broad range of the group as whole, habitat and host plant requirements are characterized by pronounced regional specialization, resulting in discontinuous and often isolated populations [1,2,3]. The taxonomy of the *Hemileuca maia-nevadensis* complex in particular has proved to be one of the most difficult among North American Lepidoptera, despite decades of intensive study of *Hemileuca* biology. Species-level taxonomic resolution has been elusive due to incongruent variation in biology, morphology, and genetic markers [3,4]. Adults and larvae vary considerably in phenotype both locally and geographically, but diagnostic morphological traits are generally lacking. Yet, host plant and habitat ecologies are highly specialized and differ among regions and populations; concomitant with reproductive barriers, there is strong evidence that multiple species-level taxa are involved [1]. Recently, genomic molecular methods have shed some light on the situation, indicating that current taxonomic concepts do not yet adequately reflect the genomic trajectory of a complex evolutionary history [5]. Six named species are currently recognized, including *H. maia* (Drury), *H. nevadensis* Stretch, *H. peigleri* Lemaire, *H. lucina* Edwards, *H. artemis* Packard.

Particularly vexing are the midcontinental taxa that occur from the Great Plains to the Great Lakes region. Great Plains populations of the *Hemileuca maia*-group have invariably been assigned to *H. nevadensis* for the last five decades (Fig 1), while the Great Lakes populations have either been assigned to *H. nevadensis* [6], or remained unassigned at the nominal species level [1, 2, 3]. Of note among the Great Lakes populations is the bog buckmoth (BBM; also called the bogbean buckmoth), its global occurrence restricted to a few isolated populations in fen habitats of the eastern Lake Ontario region, and utilizing *Menyanthes trifoliata* L. as a larval host plant. Although the BBM is of global conservation concern, its species-level taxonomy is in dispute and has variously been assigned to *H. nevadensis, H. lucina, H. maia*, unknown species status, or as a new species [3,6,7,8,9], not only hampering optimal conservation management but an understanding of the evolutionary history of the group. Such taxonomic uncertainty among the Great Lakes populations is due in part to partially shared ecological and life history traits and perceived phenotypic intergradation between named taxa. For example, adult phenotypes of Great Lakes populations appears to intergrade to the Great Plains phenotype (previously assigned to *H. nevadensis*) to the west, and to the *H. maia* phenotype to the south and east; yet differing life histories indicate that at least some Great Lakes populations represent a species distinct from not only *H. maia* and *H. nevadensis*, but all others in the group. The first molecular studies of the *maia*-group revealed surprisingly low genetic variation at odds with the accepted taxonomy [3,4], but genomic-wide molecular data has increased phylogenetic resolution, revealing that the BBM is one of the three most divergent lineages among all North American *H. maia*-group taxa 5,10].

**Figure 1.**
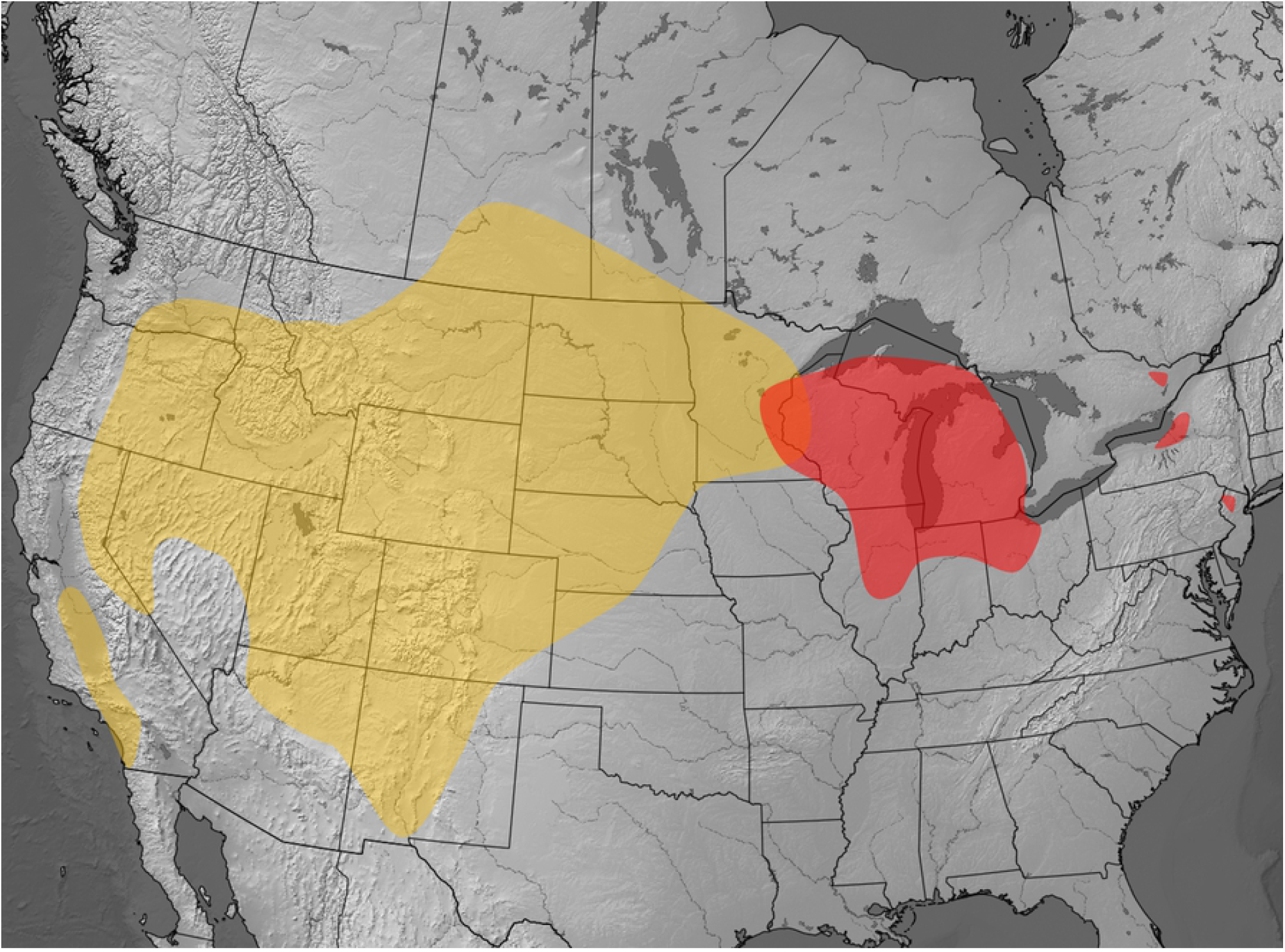
Previous concepts of the geographic distribution of *Hemileuca nevadensis*. Range of *Hemileuca nevadensis* (orange) and the *Hemileuca* Great Lakes complex (red), based on [1] and [11].

A key gap in understanding the taxonomy and biogeography of *Hemileuca* is the relationship among Great Lakes - Great Plains populations. Do they share a common post-glacial origin, do they constitute a single species, and are they conspecific with *H. nevadensis*? To begin to address these questions, *a priori* assumptions about existing taxon circumscriptions require confirmation. To this end, detailed distribution maps of all western and northern *H. maia* complex occurrences were generated, combined with analysis of genetic variation using mtDNA barcodes. Lastly, existing information on reproductive cross-compatibility between members of the *H. maia-*group was summarized, with the addition of novel data on pheromone attractiveness trials in a population of the BBM. The nomenclature of the *H. nevadensis* group is revised based on these results.

## Materials and Methods

Geographic distributions of *H. maia*-group populations from the Great Lakes westward were generated by compiling locality data from the literature [1, 2, 4, 5, 11, 12, 13, 14, 15, 16, 17], online databases [19,20] and citizen-science platforms [20, 21], with particular scrutiny of range-edge records. Occurrence dot-maps were generated with SimpleMappr [22], based on co-ordinates associated with the original data or as determined using Google Earth [23]. Occurrences precise to the county-level were mapped using approximate county-centroid coordinates. Habitat information was compiled from the literature, and *Hemileuca* site visits by the author in Alberta, Saskatchewan, Manitoba, Ontario and New York between 2001 and 2017.

Molecular variation of *Hemileuca* species was assessed using the COI “barcode” fragment (654 bp), with DNA extraction, PCR amplification, and sequencing performed at the Canadian Centre for DNA Barcoding, following standard protocols [57]. DNA sequences were analyzed on the Barcode of Life Data Systems website [46]. GLGP (Great Lakes and Great Plains) populations were sampled across the range, and all named taxa in the *maia*-group were represented. Fifty-three specimens from 23 sites were analyzed, together with 112 previously published COI sequences [3]. Voucher specimens used in a study of clinal phenotypic variation [24], deposited in the Canadian National Collection (Ottawa, Canada), were also sequenced. Voucher specimen data are given in Appendix 1. Haplotype networks were constructed with TCS 1.21, using a 95% confidence limit for parsimony [25]. The resulting haplotype network was constructed using tcsBU [26]. Haplotype variation was also visualized using a minimum spanning network, generated with the program popART [27]. To explore genetic similarity unbiased by current taxonomy, sequence cluster analysis was carried out using the BOLD interface. This analysis assigns OTU’s (operational taxonomic units) using the Refined Single Linkage algorithm [28].

Pheromone trials to test the female pheromone attractiveness of *H. lucina* and *H. maia* to male BBM were conducted in a population of the BBM near White Lake, Renfrew County, Ontario (45.30°N 76.52°W). Three live, unmated females (reared *ex* larva) of *H. lucina* from Keene, New Hampshire were placed in a 15 cm x 25 cm expandable mesh cylinder and placed in sites with actively flying males in October 2015. In September 2016, male response of BBM to *H. maia* synthetic female pheromone loaded onto rubber septa [29], placed at heights between 30 cm -100 cm on emergent vegetation.

## Results

### Taxonomy

To provide clarity and continuity in the subsequent discussion, the taxonomic changes proposed herein are addressed first, and supporting data are discussed further in the sections that follow. *Hemileuca latifascia* Barnes & McDunnough **stat. nov**. is treated as a species-level taxon for the Great Plains populations previously called *H. nevadensis*, based on the geographic isolation and genetic divergence between the two. Described from Aweme, Manitoba and previously considered a junior subjective synonym of *H. nevadensis, H*. l*atifascia* is currently the only nominal taxon described from Great Plains populations. The populations from Alberta to westernmost Minnesota and the Dakotas form the core range of *H. latifascia*; those from Nebraska to northeastern New Mexico (Fig 2) are also provisionally assigned to *H. latifascia* based on shared adult phenotype, mtDNA similarity, and shared host plant / habitat requirements. With the exception of eastern Lake Ontario discussed below, the Great Lakes populations are maintained as unresolved at the species level, pending integration of genomic and pheromone data from other key populations, particularly *H. latifascia* and the bog buckmoth. The bog buckmoth, *Hemileuca menyanthevora* Pavulaan **stat. nov**. is recognized as a species-level taxon based on the well-established unique ecology of this taxon, coupled with the recent results of [5,10] who demonstrated that the BBM is one of the three most divergent genetic clusters among all North American *H. maia*-group taxa (the remaining two clusters comprising nominate *H. nevadensis* + *H. artemis*, and combined *H. maia* + *H. lucina* + Great Lakes Complex, with the caveat that the latter are incompletely sampled). Application of the name *H. menyanthevora* should be limited to the eastern Lake Ontario populations, which are widely divergent from other Great Lakes complex populations sampled to date [5, 10]. Taxonomic changes were not proposed by [5, 10], and two names for the BBM, *Hemileuca maia menyanthevora* Pavulaan [8] and *Hemileuca iroquois* Cryan & Dirig [9], were published subsequently. Both names were described based on BBM populations from Oswego Co., New York. The description of *H. iroquois* [9] was published five days after that of *menyanthevora* [8], and *H. iroquois* Cryan & Dirig is therefore revised to junior synonymy (**syn. nov**.) following the ICZN guidelines for publication priority [30]. In summary, GLGP *Hemileuca* populations are divided into *H. latifascia* of the Great Plains, *H. menyanthevora* of eastern Lake Ontario, and intervening Great Lakes / Upper Midwest populations are maintained as unresolved at the species level (Fig 2).

**Figure 2.**
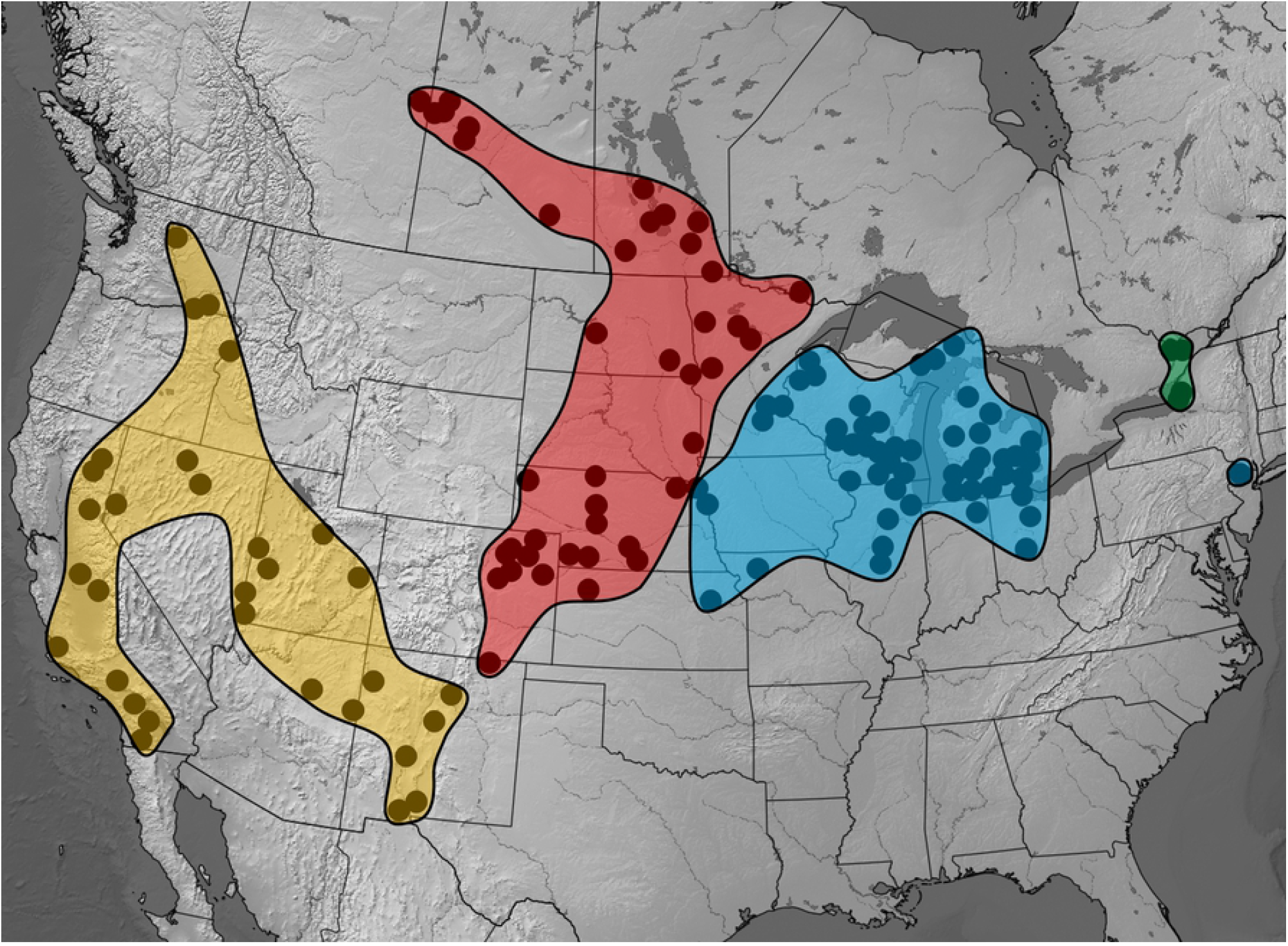
Revised geographic distribution of *Hemileuca nevadensis* and related species. Revised range of *Hemileuca nevadensis* (orange), *Hemileuca latifascia* (red) and “Great Lakes Complex” (blue) as determined in this study; *Hemileuca menyanthevora* (green).

### Geographic distribution and habitat

Despite previous depictions of distribution spanning the entire Rocky Mountain-western Plains region (Fig 1), combined historic and modern occurrence records of *Hemileuca nevadensis* reveal a vast region centered on Montana-Wyoming where no occurrences of *H. nevadensis* (or any other *H. maia*-group species) exist (Fig 2). The range of *H. nevadensis* has previously been depicted as extending from California to the central and northern Great Plains (e.g. [1, 3, 9, 11]). The closest occurrences between the Great Plains and those to the west are in New Mexico, although even these populations are separated by about 150 km and a major mountain range: a Colfax County population cited by [11] at the western edge of the Great Plains, separated by the Sangre de Cristo Mountains from the northernmost occurrences in the Rio Grande drainage of Santa Fe County [21].

The northern range limits comprise ecologically similar but very discrete, geographically isolated populations restricted to dune fields of the Aspen Parkland ecoregion in eastern Alberta, southern Saskatchewan and southwestern Manitoba. Those of the Edgerton dune field in eastern Alberta [16, 31] form the northwestern limit (Fig 2). No *Hemileuca maia*-group records are known from Montana, Wyoming, southern Alberta, southwestern Saskatchewan, the western Dakotas (notably including the entomologically well-studied Black Hills, South Dakota), eastern Idaho or British Columbia (*contra* [1, 3]; see [32]). Other minor differences in range limits are attributed to the recent (post-1996) discoveries of populations that slightly extend the known range, including Rainy River, Ontario; Monona County, Iowa; and Riley County, Kansas.

The northern Great Plains populations (Alberta, Saskatchewan, Manitoba) are closely tied to dune habitat characterized by a stunted growth-form of trembling aspen (*Populus tremuloides* Michx.), a plant community moderated by the edaphic conditions of extensive aeolian dune fields [31]. These sites are situated in the parkland ecoregion, whereas geologically similar dune fields further south in the prairie ecoregion lack *Hemileuca* (despite intensive collecting effort in the last two decades; see e.g. [16]), presumably due to the scarcity or absence of the aspen stands that are present in more northern dune fields. Populations in Colorado and central Nebraska are also associated with major dune fields, but larvae instead feed on willows growing along waterways. The association with sand may be linked to the pupal stage, as pupation occurs on or underneath the soil layer [1].

### mtDNA variation

Nominate *H. nevadensis* from Nevada and California is represented by three unique haplotypes with a minimum divergence of 2.6 % (uncorrected pairwise distance) from all other North American *maia*-group taxa. These formed a separate cluster unlinked to the remaining haplotypes in the network parsimony analysis (Fig 3). The sequence cluster analysis segregated all *maia*-group sequences into two OTU’s, California-Nevada *H. nevadensis* versus all remaining samples. GLGP populations from Alberta, Manitoba, northern Michigan, Wisconsin and Ontario exhibited a single haplotype (haplotype *1a*; Fig. 4), with two similar haplotypes from Roscommon County in central Michigan (haplotypes *1c, 1d*; Fig 3), and another shared between New York *H. menyanthevora* and Massachusetts *H. lucina* (*1b*; Fig 3). The willow-feeding populations from southern Michigan and northern Ohio that have adults similar to *H. maia* shared a haplotype that occurred westward into Iowa, Nebraska and Colorado (haplotype *2a*; Fig 3). Northern *H. maia* samples and *H. peigleri* were similar to this latter cluster of GLGP populations, whereas *H. slosseri* and *H. artemis* formed a separate group more similar to the northern *H. latifasia* haplotypes and southeastern *H. maia*. Florida *H. maia* was quite divergent by comparison, even from nearby populations in Georgia. The GLGP haplotypes *3* and *4* were unique to Wisconsin and Nebraska, respectively.

**Figure 3.**
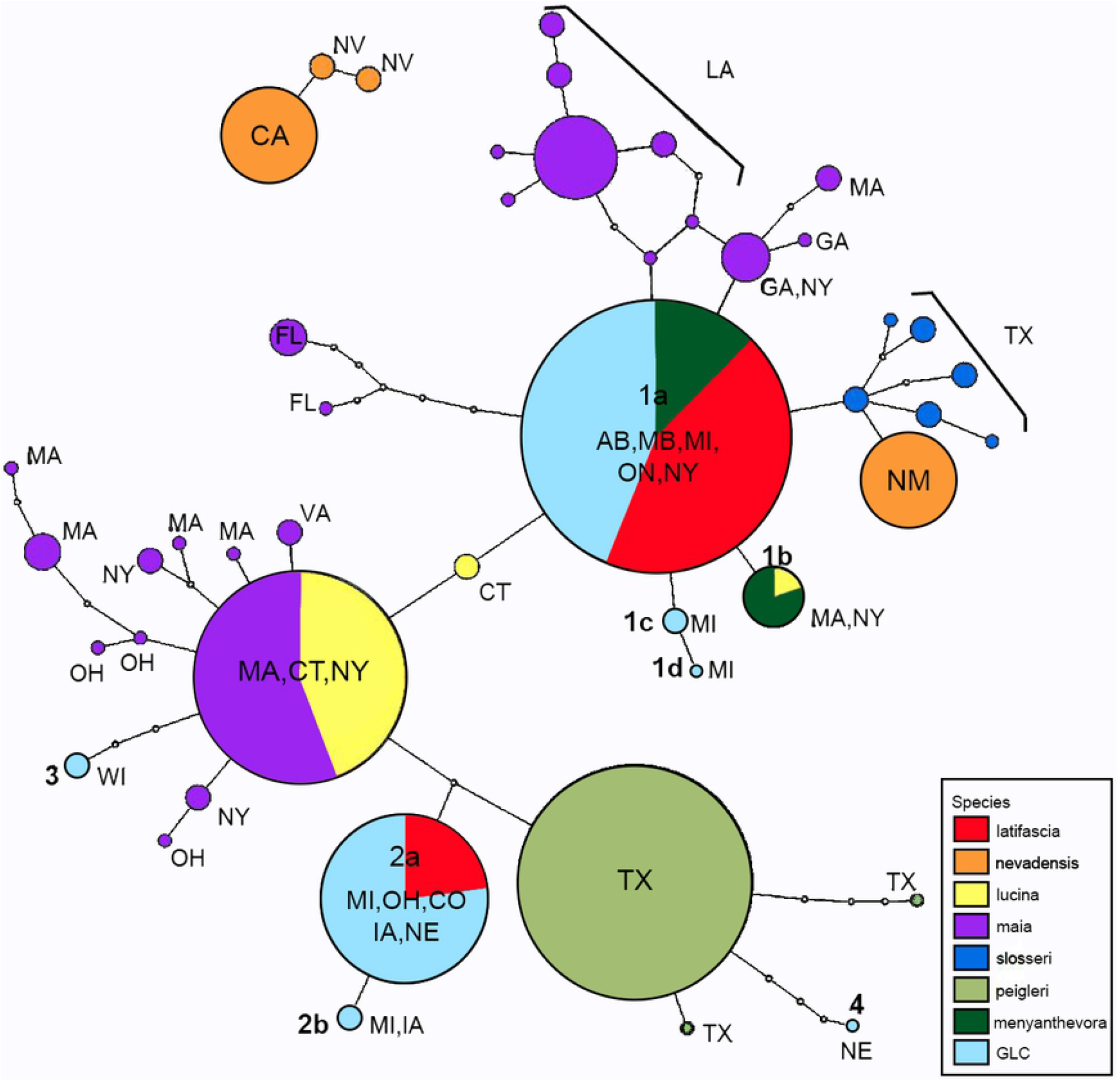
*Hemileuca maia*-group haplotype network. *Hemileuca* haplotypes (95% confidence limits; [25]) based on the COI-barcode sequence (654 base-pairs). Colored circles represent observed haplotypes, size is proportional to number of samples and color corresponding to species. Haplotypes of H. latifascia, H. menyanthevora and Great Lakes Complex (GLC) are numbered 1a-1d, 2a-b, 3, and 4. Two-letter state/provincial abbreviations indicate sample origin.

**Figure 4.**
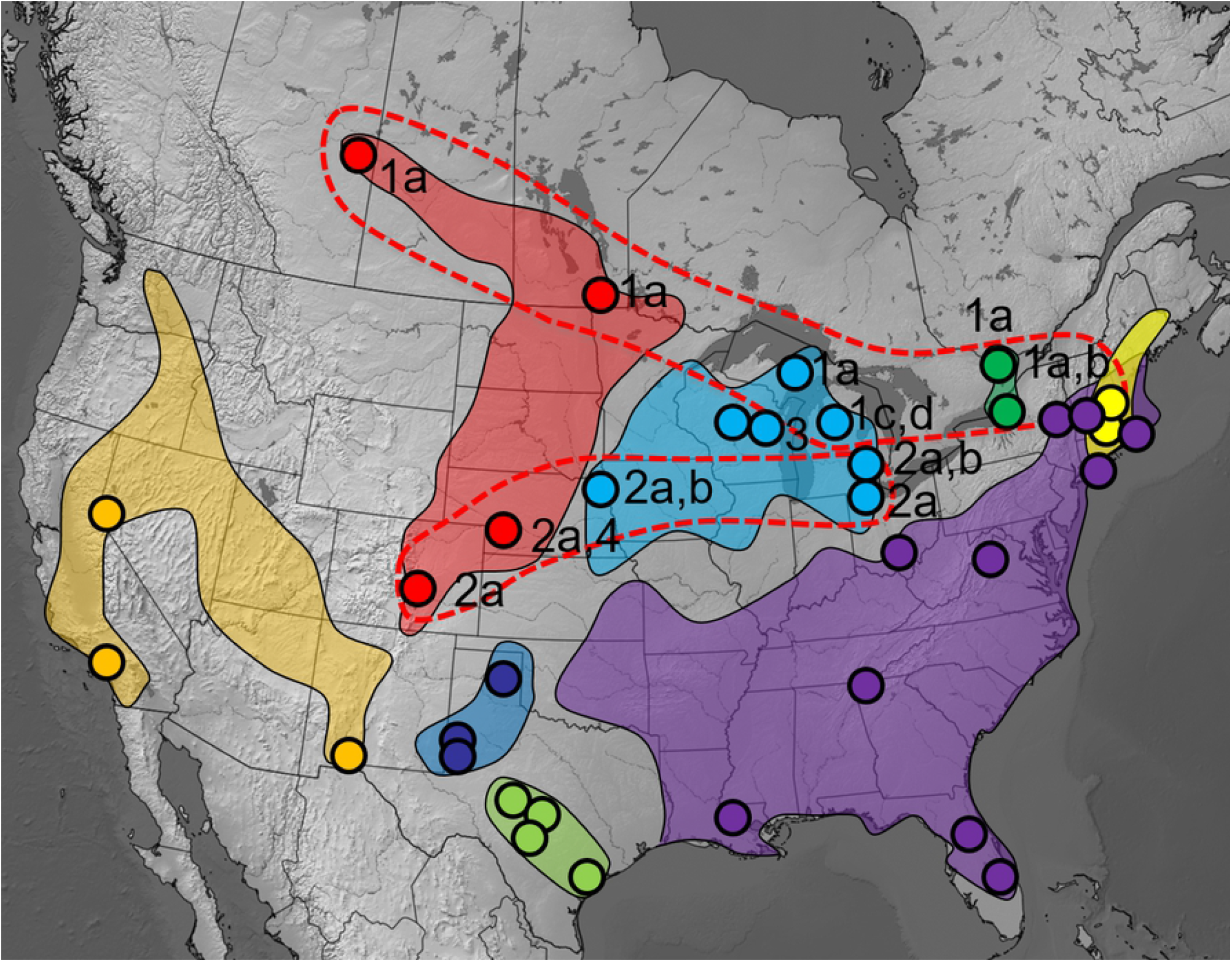
Geographic range and origin of *Hemileuca* mtDNA haplotypes. Taxon range (shaded) and sample origin (circles) of *Hemileuca* mtDNA vouchers, with corresponding haplotype numbers from Figure 3. *H. nevadensis* (orange), H. latifascia (red), Great Lakes Complex (light blue), *H. menyanthevora* (dark green), *H. slosseri* (dark blue), *H. peigleri* (light green), *H. maia* (purple), *H. lucina* (yellow).

### Pheromone trials

Synthetic pheromone lures of *H. maia* failed to elicit a response from *H. menyanthevora* males. Approximately 20 male *H. menyanthevora* were observed flying in the core habitat at the White Lake site while lures were deployed, but no males approached the lures. Virgin female *H. lucina* also failed to attract male *H. menyanthevora* in the previous season (2015). At least 19 pheromone cross-trials have been documented for GLGP populations (Fig 5; Table 1), including comparisons to *H. maia, H. lucina, H. artemis* and among populations of GLGP *Hemileuca*.

**Figure 5.**
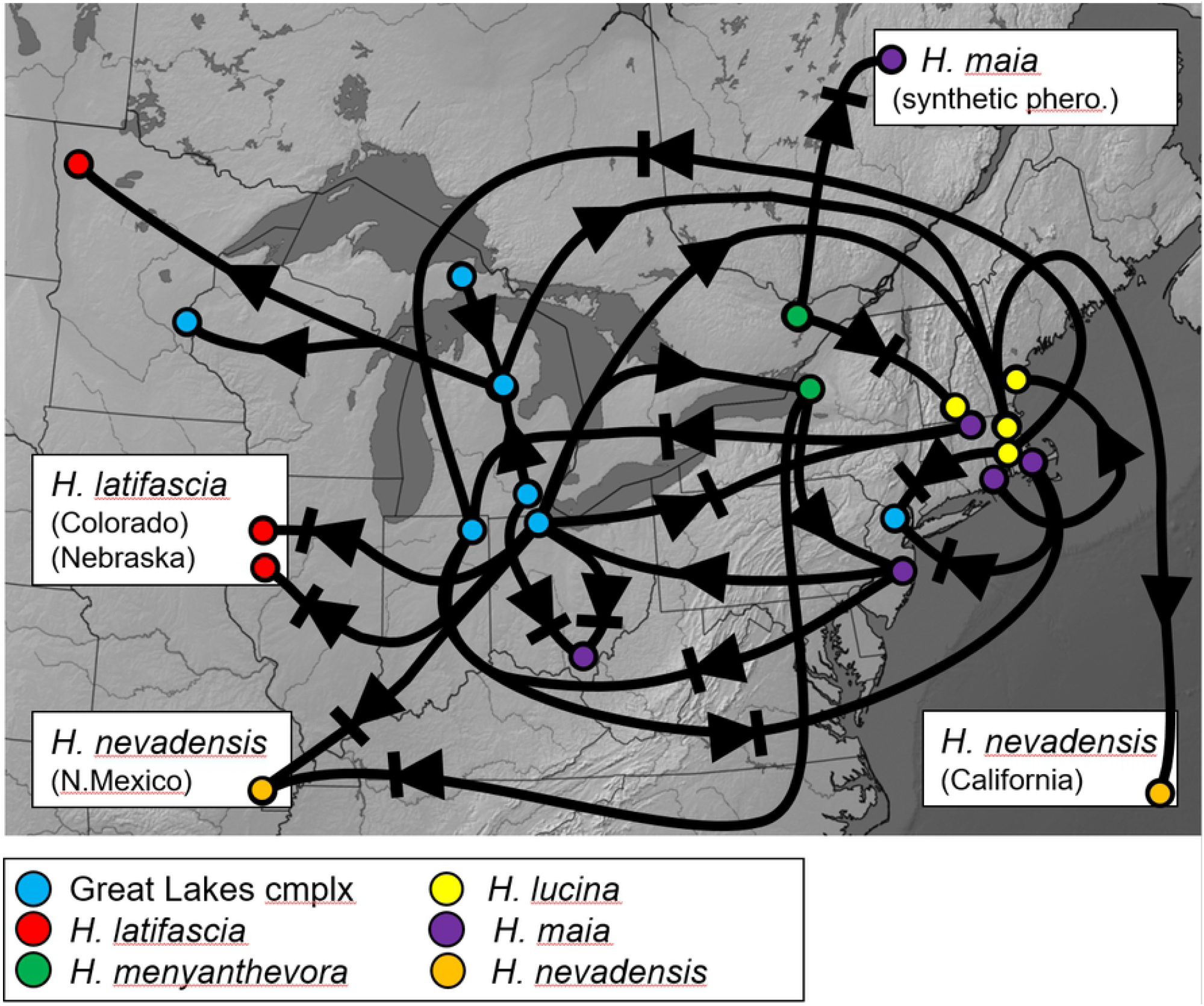
Geo-referenced summary of *Hemileuca* pheromone cross-attraction trials. Circles indicate geographic origin of tested individual, direction of arrow indicates male-to-female attraction, barred arrow indicates lack thereof. Based on the results summarized in [1, 24] and this study.

**Table 1.** Summary of female pheromone trials involving *Hemileuca latifascia*. Results based on [1, 24], and this study.

| Cross | Number of cases |  |
| --- | --- | --- |
|  | Positive male response | No male response |
| *latifascia* - *maia*<sup>1</sup> | 2 | 7 |
| *latifascia* - *lucina* | 1 | 3 |
| *latifascia* - *latifascia* | 3 | 1 |
| *latifascia* - *artemis* | 0 | 2 |

<sup>1</sup> includes synthetic pheromone lure results

## Discussion

### Distribution, habitat and host plants

*Hemileuca nevadensis* was thought to occupy a large, contiguous range spanning from coastal California across the Rocky Mountain region to the Midwest [1, 3, 4, 7, 9, 11], but there are in fact no populations known for the vast area between the Canadian Prairie Provinces and the Great Basin. This gap covers over 1.3 million km^2^, or approximately 30% of the presumed range. What has been called *Hemileuca nevadensis* therefore actually consists of two allopatric ranges with a wide distribution gap across the Rocky Mountains and western Great Plains. In the 50 years since Ferguson’s landmark Saturniidae monograph [11] that sparked intensive study of the group, no populations have been discovered in this region, including all of Montana and Wyoming (Fig 2). Lemaire cites an unspecified Montana record [2], but without further data. It is possible (and indeed likely) that undiscovered *Hemileuca* populations exist in eastern Wyoming and Montana – the region is one of the most undersampled for Lepidoptera. It is however unlikely that undiscovered populations linking Great Plains – Great Basin occurrences across the Rocky Mountains, as this region has received considerable survey effort for over a century. None of the recently-discovered populations are within this large gap, instead being relatively close to known populations.

Like *H. nevadensis, H. latifascia* uses primarily Salicaceae as larval hosts, but unlike many Saturniidae, host plant is not a limiting factor in GLGP *Hemileuca* geographic distribution. Prairie Province populations are highly localized to dunes, yet the trembling aspen and willow that serve as larval hosts are the most abundant tree and shrub cover across the parkland region [33]. In the Great Lakes region, a variety of unrelated deciduous wetland shrubs comprise the larval hosts [17]. In Manitoba larvae sometimes occur on bur oak [12], and the Barry County, Michigan population (possibly now extinct) that fed on black oak remains a puzzle [1]. The reliance of *H. menyanthevora* on *Menyanthes* has perhaps been over-emphasized as a taxonomic trait, because larvae also feed on a wide variety of other wetland plants [34], and a Wisconsin *Menyanthes*-feeding population is not closely related to *H. menyanthevora* [5]. Therefore, *Menyanthes*-feeding as an opportunistic shift to a host plant in the right microhabitat (likely having occurred more than once) provides a better explanation of host use. Additionally, *Menyanthes* occurrence is poorly correlated with that of *H. menyanthevora*; a common peatland plant, it is found throughout the many bogs and fens of eastern Ontario, but *H. menyanthevora* is restricted to only two fen complexes [7]. *Menyanthes* phenology also restricts herbivory by *H. menyanthevora* in that spring leaf-out sometimes occurs well after larvae have hatched, forcing use of alternate hosts. However, the ability for *H. menyanthevora* larvae to develop entirely on this plant appears to represent a unique evolutionary adaptation, as *Salix*-feeding populations from Michigan cannot develop successfully on it [13]. Northern *H. latifascia* populations underwent a shift to *Populus tremuloides* on dunes (either subsequent to or independent of the switch to wetlands), permitting a northwestward expansion. GLGP *Hemileuca* mostly occur in habitats with an abundance of *Salix*, but a switch to non-*Salix* hosts is evident where this shrub is (or became) too infrequent to sustain *Hemileuca* populations. In the eastern lake Ontario region this switch may have occurred relatively recently in response to postglacial plant community changes.

Larval hostplant does not limit broad-scale distribution of GLGP *Hemileuca*, which is instead determined by habitat availability. Range-edge habitats consist of sand barrens, dunes and treeless wetlands associated with landscape features such as lakeshores, glaciolacustrine and glaciofluvial surface geology. About 75% of the modern-day GLGP range was ice-covered during the last glacial maximum [35], and most northern populations are associated with sand deposits, particularly those associated with the southwestern shoreline of glacial Lake Agassiz [36]. Northern Minnesota and southeastern Manitoba populations are on the former lake-bed of glacial Lake Agassiz, which reached its maximum extent approximately 11 ka (thousand years ago) [36]. Eastern Ontario populations are within the limits of the Champlain Sea, a temporary incursion of Atlantic waters that existed from about 13 – 10 ka. No populations occur on the Canadian Shield (the White Lake, Ontario population is on the edge of the Shield but is underlain by marble), and despite the abundance of wetlands the geology and topography of the Canadian Shield are generally not conducive to the development of suitable habitat.

### Genetic variation

Compared to the four nuclear markers employed in previous studies, mtDNA provides better resolution among *Hemileuca maia* complex taxa, yet does not recover all taxa as monophyletic groups [3]. *Hemileuca nevadensis* from California and Nevada is genetically the most divergent of all *maia*-group species [5], even with expansion of geographic sampling to virtually all regions; haplotype parsimony analysis did not resolve haplotypes of nominate *H. nevadensis* as connecting to any remaining *H. maia*-group populations (Fig 3). At 2.6 % minimum divergence, *H. nevadensis* is somewhat above the average minimum of 1.5 - 2 % interspecific divergence for other North American Lepidoptera, consistent with the interpretation that nominate *H. nevadensis* is a species distinct from all others in the group. With a divergence estimate of about 4% based on a longer mtDNA sequence (∼1500 bp; [3]), the time of divergence between *H. nevadensis* and Great Plains populations dates to approximately 2 mya (assuming 1 % divergence per lineage per million years; [37]). This approximates the onset of the Quaternary glaciations at 2.4 mya [38], and would imply that the genetic variation in the remainder of the *maia*-group was largely shaped by the palaeo-ecological changes of the Quaternary period over the last 2 million years. Phylogeographic data unambiguously delineate traditional *H*. “*nevadensis*” as actually consisting of two parapatric, divergent taxa: nominate *H. nevadensis* west of the Rocky Mountains, and *H. latifascia* of the Great Plains – Great Lakes. Other Great Basin and especially New Mexico - Arizona *H. nevadensis* should be assessed, and they are here provisionally aligned with nominate *H. nevadensis* based on phenotypic similarity and phylogeography of other Lepidoptera in western North America [39].

The remaining *maia-*group populations exhibit an overall pattern of limited genetic similarity within geographic regions rather than discrete species clusters [3], but the addition of northern populations shows a striking deviation from this pattern in that a single haplotype is not only seemingly prevalent within several populations, but also has the widest distribution of any *maia*-group haplotype (Fig 4). There is no indication of a major east-west phylogeographic break between Great Lakes populations and those previously depicted as “*nevadensis*” in the Great Plains (Fig 4). On the contrary, there is greater genetic similarity between populations spanning from Alberta to northern New York, and southern Michigan to Colorado, than between any other geographic segregates. This latitudinal stratification is consistent with a vast postglacial expansion into the east-west periglacial habitats that formed after the recession of the continental ice sheet.

*Hemileuca menyanthevora* samples have the same or similar haplotype as other northern GLGP populations, consistent with a genetic signature expected from post-glacial expansion due to founder effects, and possibly also introgression in the case of haplotype sharing between *H. menyanthevora* + *H. lucina* and *H. menyanthevora* + *H. latifascia* (Fig 4). Despite the low taxonomic resolution of the COI data, haplotype-sharing between species is rare: using a longer COI sequence, Rubinoff et al. (figure 3) illustrate two instances of sharing [3], one between Texas populations of *H. peigleri* and *H. maia* an the second between Nebraska *H. nevadensis* and Iowa *H. maia*. The *maia* + *peigleri* example appears to be a labelling error since no Texas *H. maia* were sampled in that study, and those samples seem instead to refer to the recently-discovered populations on the Texas coast, identified elsewhere in that paper as *H. peigleri*. The Iowa population referred to *H. maia* in that study is here classified as *H. latifascia*, as the host plant and distinctive larval phenotype of that population [40] makes identification as a dark phenotype of *H. latifascia* more tenable, as does geographic location and the COI haplotype shared with *H. latifascia* populations from four other states (Fig 3).

The mitogenomic variation of the Michigan-transect populations analyzed by [24] is unexpected, and provides some unique clues to the origin and history of the Great Lakes *Hemileuca*. Adult phenotypes of this transect show a transition from translucent wings in northern populations to darker and less translucent populations that are more *maia*-like in appearance in southern Michigan, northern Ohio, and possibly as far west as Iowa in a recently-discovered population in the Loess Hills. It was previously postulated that this pattern indicated clinal, north-south variation in adult phenotypes (from *lucina*-like to *maia*-like), representing meta-populations of the same species. Upper Peninsula (Schoolcraft Co.) samples shared a geographically very widespread haplotype (*1a*; Fig 4), from Alberta to Ontario and New York, and similar central Michigan haplotypes. Yet, southern Michigan and northern Ohio samples are imbedded in a different part of the haplotype network, closer to *H. maia* haplotypes (Fig 3). The presence of two genetic groups instead suggests separate founding populations across Michigan. The group 2 haplotypes of *H. latifascia* may represent past contact with *H. maia* as previously suggested [24], but even geographically proximate populations of *H. latifascia* – *H. maia* are now reproductively isolated (Fig 5). Geographic occurrence of *maia*-like *latifascia* gives further evidence of potential historical contact, as *maia*-like populations are found near the southern periphery of *H. latifascia*. What was initially thought to be north-south clinal variation in *H. latifascia* instead appears to represent a complex population history of at least two lineages. The fact that *H. maia* and *H. latifascia* are reproductively isolated would indicate that genetic exchange is not recent, or that isolating mechanisms including pheromonal incompatibility evolved very rapidly. Lastly, evidence from host plant performance trials also points to greater similarity of southern Michigan populations to *H. maia* than *H. latifascia* further north: *maia*-like *H. latifascia* from Lucas Co., Ohio showed similar survivorship to oak-specialist *H. maia* when reared on *Quercus rubra*, (57 and 62 % respectively, first 12 d after hatching), while *H. latifascia* from New York experienced only 1% survival on this host [13].

*Hemileuca maia* exhibits several haplotype clusters in separate parts of the haplotype network. Multiple glacial refugia may have been involved in shaping this genetic diversity, and different clusters for Appalachian, southern coastal, and Floridian populations are evident. Regional differentiation may be ongoing, and more intense geographic sampling of *H. maia* would provide more precise measures of apparent phylogeographic structuring. *Hemileuca peigleri* has been treated as a subspecies of *H. maia*, and indeed the mtDNA data indicate a closer relationship to *H. maia* than *H. latifascia*, consistent with previous results [3]. Conversely, the Texan species *H. slosseri* is more similar to *H. artemis* and *H. latifascia* than *H. maia* or *H. peigleri*.

### Pheromone cross-compatibility

Pheromone compatibility presents a particularly valuable taxonomic character set in the *Hemileuca maia*-group, as mating barriers exist between most of the taxa currently recognized as species. this is in contrast to other well-studied saturniid groups that lack such barriers (e.g. *Hyalophora, Automeris*; summarized in [1].

Seven of nine trials between GLGP *Hemileuca* vs. *H. maia* indicate that female pheromone is not cross-attractive, including geographically adjacent populations in the eastern Lake Erie region (Fig 5; Table 1), and *H. maia* synthetic pheromone elicits no response in *H. menyanthevora*. Sympatric populations of GLC and *H. maia* in New Jersey also remain distinct (NatureServe 2017), and pheromones are not cross-attractive [41]. There is therefore good evidence that strong prezygotic reproductive barriers exist.

Of the four cases testing pheromone cross-attraction between *H. latifasca* versus *H. lucina*, three produced negative results (Fig 5). Massachusetts *H. lucina* females can reportedly attract *H. latifascia* males from Lucas Co., Ohio and produce viable F1 females [1], and this cross warrants further scrutiny. Evidence of pheromone non-compatibility between *latifascia* and *lucina* is therefore somewhat more tenuous than *latifascia*-*maia* due to lower sample size, but taken together with the phenotypic and ecological differences of parapatric *latifascia* and *lucina* populations in northeastern USA, and the absence of the widespread group-*1* haplotypes in *lucina*, the two should be maintained as separate species-level taxa.

Cross-attraction of *latifascia* and *artemis* has been tested twice, and in neither instance was a positive male response observed; this cross can however produce viable F1 hybrid females [1]. Lemaire [2] and Rubinoff et al. [3] treat *artemis* as a species distinct from *nevadensis*, with which it was previously synonymized [1]. Genomic data do not support recognition of *H. artemis* as a species distinct from *H. nevadensis* [5]. Evidence for reproductive isolation between *latifascia* and *artemis* as indicated by pheromonal and genetic distinctness and geographic isolation, further support that *artemis* is not closely related to *H. latifascia*. No pheromone trial data are available for *latifascia* vs. *nevadensis*.

A single trial comparing *latifascia* between Great Lakes and Great Plains populations did not elicit a positive response: Lucas County, Ohio males were not attracted to northeastern Colorado females, but three other interpopulation trials among *latifascia* elicited a positive response ([1]; Fig. 5). Further Great Lakes vs. Great Plains cross-trials should be attempted to determine if the apparent lack of response between Ohio-Colorado holds true elsewhere.

Some geographically distant and unrelated populations can exhibit pheromonal compatibility (but result in infertile F1 hybrids), such as California *H. nevadensis* versus New England *H. lucina*, (Fig. 5). However, interpretation of pheromone results must be considered in the light of the fact that there is little or no selective pressure for widely allopatric populations to maintain reproductive isolating barriers [42]. Intraspecific geographic variation in pheromone components may also play a role (e.g., [43]).

### Taxonomy

Great Plains populations of the *maia-*group were originally described as a subspecies, i.e. *Hemileuca lucina latifascia* [44], the only taxon named from the Great Plains – Great Lakes region (Type locality: Aweme, Manitoba). As more information on *Hemileuca* populations came to light over the next 50 years, Ferguson hypothesized that these Great Plains/Great Lakes populations were instead conspecific with *H. nevadensis*, arguing that *H. nevadensis* was a phenotypically variable species with a distribution spanning from coastal California east to Wisconsin and Illinois [11]. This has essentially remained the taxonomic *status quo* for *H. nevadensis*, with differences in species designations for the problematic Great Lakes populations ranging from *H. nevadensis, H. maia*, hybrids thereof, *H. lucina* to simply unknown (e.g. [2, 24]). The Great Plains populations are now generally recognized as being conspecific with *H. nevadensis* based on ecology and reproductive isolation from *H. maia*, but exhibiting *H. maia*-like phenotypes in some populations [6]. This previous delineation of *Hemileuca nevadensis* is not congruent with the molecular, biogeographical and ecological data summarized here. Northern Great Plains – Great Lakes populations are genetically similar, whereas nominate *nevadensis* is the most divergent among the whole group. The two groups have allopatric ranges that are separated by a vast distributional gap consisting of most of the Rocky Mountain region. The genetic distinctness of nominate *H. nevadensis* in parallel with a redefined, allopatric distribution provides compelling evidence that Great Plains – Great Lakes populations are a biogeographically and genetically cohesive group that is not only distinct from true *H. nevadensis*, but that *H. nevadensis* may in fact be the sister-species to the rest of the *maia*-group [3]. Adult phenotypic traits unique to nominate *H. nevadensis* not shared by Great Plains populations include thoracic coloration, wing discal spot size and forewing shape [11, 44]; diagnostic adult traits are essentially lacking in all other *maia*-group species.

Although some GLGP populations were historically sometimes associated with the taxon *H. lucina* at the species level, the latter is now widely accepted as being endemic to New England and not conspecific with any GLGP taxa (e.g. [1, 3, 5]). The relationship of GLGP *Hemileuca* to *H. maia* is somewhat more complicated, owing to a larger region of potential contact (Fig 2) and the existence of unrecognized genomic diversity within what is currently treated as a single species [3, 5]. However, pheromone cross-attraction trials, habitat and host plant, and genomic variation provide strong support that *H. maia* is not conspecific with any GLGP populations. Although the geographic and taxonomic limits of the Great Plains *Hemileuca* taxon requires further study, it is demonstrably distinct from all other named species in the group and should therefore be recognized as *Hemileuca latifascia*. The Colfax County, New Mexico population (Ferguson 1971) is here provisionally associated with those of the Great Plains, following [2], but given that the nearest *H. latifascia* populations in eastern Colorado occur in a different ecoregion and habitat, the association of this population with *artemis* or even nominate *nevadensis* should be evaluated.

The adult and larval variation of *Hemileuca latifascia* and the Great Lakes populations are detailed by Tuskes et al. [1], and an apparent clinal zone exists in northwestern Wisconsin, where adults can have wide, narrow or intermediate medial bands within the same population [45]. Larvae of *H. latifascia* vary from mostly yellow, to almost entirely dark in eastern Manitoba [12], while Alberta populations are mostly of the striped-yellow form (Fig 6) with yellow phenotypes rare. The *maia*-like populations from the southern range portions represent another distinct phenotypic group, although larvae and ecology are mostly like that of other Great Lakes populations. A comparison of genetic and ecological differences among the groups recognized herein is given in Table 2.

**Figure 6.**
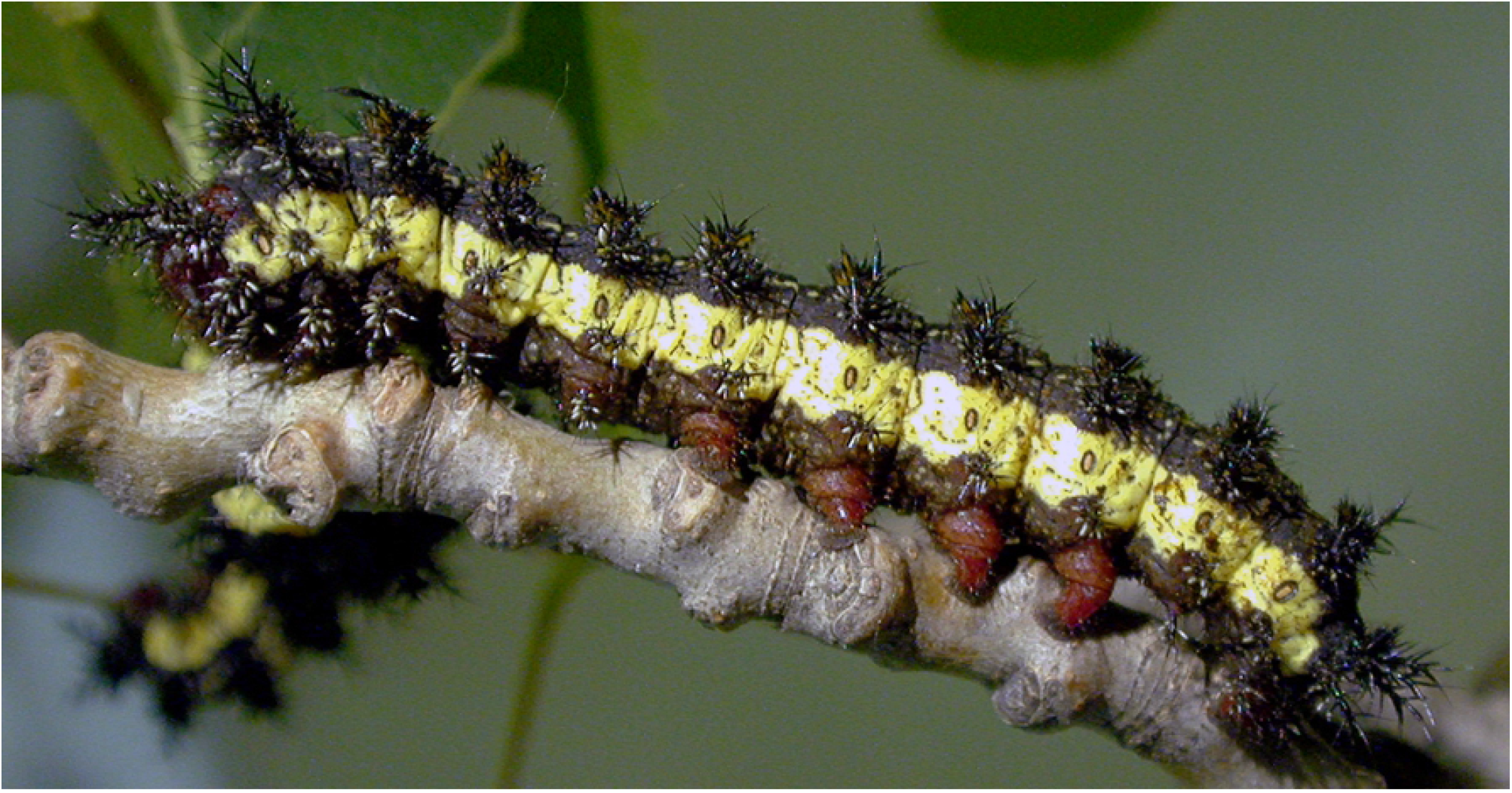
*Hemileuca latifascia* larva. Edgerton dunes, near Edgerton, Alberta, Canada. C. Schmidt photo.

**Table 2.**
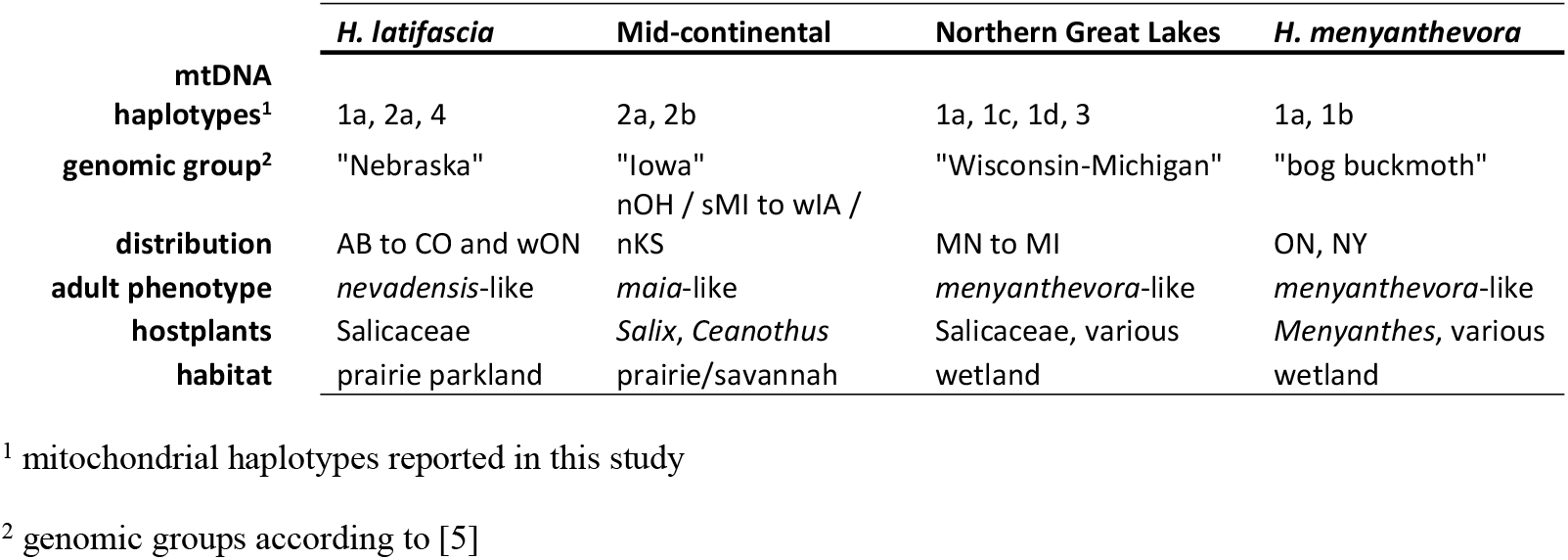
Comparison of genetic and ecological differences among Hemileuca latifascia and related taxa.

### Glacial refugia and post-glacial expansion

The *Hemileuca* populations that are taxonomically most problematic occur in or near regions that were ice-covered during the last glacial maximum [47], which occurred about 20 ka [35]. Understanding potential glacial refugia, postglacial dispersal routes and contact zones would therefore aid taxonomic hypotheses. Many LGM plant communities bore little resemblance to those of today [48], but the most plausible refugia for GLGP *Hemileuca* would have been warm-temperate non-forest habitats with abundant Salicaceae, which persisted in the southern Great Plains region as grassland throughout the LGM [49]. The most parsimonious hypothesis is that riparian corridors and dune fields similar to those inhabited on the Great Plains today served as a south-central Great Plains refugium for *H. latifascia*. Two geographic clusters are embedded within the haplotype network, the northern group (haplotypes *1a,1b,1c*) from Alberta to New York, and the southern group (*2a,2b*) from southern Michigan through the central Great Plains. This latitudinal stratification could be the result of separate refugia, such as a mid-continental or Mississippi valley refugium typified by a cooler and wetter biome during the LGM [48] serving as the source for the northern haplotype group, whereas a separate southwestern refugium could explain mitochondrial DNA similarity of haplotype *4* to *H. peigleri* of central Texas (Fig. 3). The shared mtDNA haplotypes between *H. menyanthevora* and northern GLGP populations, despite genomic divergence of *H. menyanthevora* (Dupuis et al. 2018, 2020), indicates recent genetic exchange among these populations, despite a greater and likely older divergence of *H. menyathevora*. Determining the evolutionary origins of *H. menyanthevora* in the context of GLGP *Hemileuca* will require integration of genomic data from key populations that link those of the Great Lakes region to *H. latifascia*: do *H. menyathevora* and *H. latifascia* share a recent common origin as mtDNA variation suggests, or is this simply a result of mtDNA introgression masking an older evolutionary divergence of *H. menyanthes* from other GLGP *Hemileuca*?

As currently defined, the global range of *H. menyanthevora* lies entirely within the maximum extent of the Wisconsinan glaciation [35]. Contrary to previous hypotheses, genomic evidence does not support recent origins of *H. menyanthevora* from any other *H. maia* group populations and its Pleistocene history remains enigmatic. The postglacial expansion of *Hemileuca menyanthovora* to its modern-day distribution would have been blocked initially by the retreat of the last ice lobes and subsequently the Champlain Sea which persisted until about 10 ka [36], followed by unsuitably cold-boreal environments maintained by anticyclonic glacial weather patterns (cold, dry easterly winds induced by the receding continental ice sheet; [50]). Despite the current lack of molecular evidence, a hypothesis favoring a midcontinental refugium for *H. menyathevora* is perhaps still most likely, inasmuch as geographic proximity across shared habitats indicate connectivity across the Great Lakes and northern Great Plains *Hemileuca* populations, as opposed to a southern Appalachian or Atlantic coastal refugium. However, the existence of a taxonomically unresolved wetland population of *Hemileuca* in northwestern New Jersey [6,47] begs further research.

Under a midcontinent refugium scenario, dispersal of *Hemileuca* into postglacial GLGP was likely correlated with expansion of prairie habitats into eastern North America (the Prairie Peninsula) about 6 ka and 8 ka [51, 52]. Conversely, the Prairie Peninsula and absence of oaks would have prevented post-glacial expansion of the oak-obligate *H. maia* until much later [41]. Colonization of postglacial habitats by GLGP *Hemileuca* would have been facilitated by the ability to use larval hosts plants other than oak, as oak had a restricted LGM range, and was slower to colonize post-glacially [48, 53].

The lower peninsula of Michigan was one of the earliest ice-free regions, and together with extensive surficial sand deposits, may have provided the first extensive suitable habitat in the Great Lakes and acted as a source region for eastward and westward expansion. The largest known populations of GLGP *Hemileuca* persist there today [17]. The drastically lower water levels of Lake Huron approximately 7 ka, exposing what is now lake bed [36], would have provided a broad dispersal route into southern Ontario, in concert with the climatic warming that saw the recession of cold-boreal plant communities from the basin of the former Champlain Sea [53]. Favourable (non-forested) *Hemileuca* habitat such as wetlands would have been more extensive prior to recession of postglacial wetlands and afforestation to modern-day biomes. The highly isolated *H. menyanthevora* populations speak to a more expansive postglacial distribution, and could be explained by colonization through the Champlain Sea basin. Subsequent climatic oscillations and concomitant plant community changes would have resulted in localized extinctions from much of the eastern range limits of *H. menyanthovora*, leaving a few highly isolated populations. The Eastern Prairie Fringed Orchid (*Platanthera leucophaea* Nutt. (Lindl.)), a threatened species of calcareous fens, occurs with *H. menyanthovora* and *H. latifascia* at several sites and has a strikingly similar distribution pattern. Of note is the New Jersey wetland *Hemileuca* population of unresolved status [6], which shares a similarly disjunct occurrence with the Eastern Prairie Fringed Orchid in the marble belt of northern New Jersey [54]. If indeed related to *H. menyathevora*, it would indicate postglacial dispersal links between widely separated regions, either through the Richelieu Valley of Quebec / Vermont or from eastern Lake Ontario through the Mohawk Valley of New York. Alternatively, the New Jersey taxon may be derived from the mid-continental *maia*-like group, the nearest of which persist in northwestern Ohio. This would imply a postglacial colonization across the region south of Lake Erie and Lake Ontario, which at one time was home to many wetlands that were drained as early as the late 1800’s [55]. The likelihood of a shared postglacial history of GLC *Hemileuca* on either side of the Appalachian divide warrants further investigation.

*Hemileuca latifascia* populations of the Canadian Prairie Provinces more plausibly represent an eastern expansion from upper Midwest *Hemileuca* populations, rather than a northward expansion across the western Great Plains, given the dissimilarity between Alberta-Manitoba versus Colorado mtDNA haplotypes, the much greater distribution gap between Alberta vs. Colorado populations than Alberta vs. Manitoba, and the unique trembling aspen/dune habitat shared across Prairie Province sites. The Holocene vegetation history of large dune fields in central Saskatchewan indicate that suitable habitat existed there approximately 5 ka [56], a location that is situated as a midway stepping stone between modern-day *H. latifascia* populations in eastern Alberta and western Manitoba.

A similarly heterogeneous haplotype complement for *Hemileuca maia* probably also reflects the genetic legacy of multiple glacial refugia, and many populations are regionally isolated still today. Palynology and biome reconstruction for the LGM provide somewhat more confidence in refugial hypotheses for *H. maia* since *Quercus* pollen is one of the most important components in extrapolation of palaeo-environments. *Quercus* was limited to the extreme southern United States during the LGM [49] so the *Quercus*-obligate, thermophilic *H. maia* must have had a similarly restricted range. Midcontinental / Appalachian, Gulf Coastal, and Floridian haplotype clusters are consistent with the presence of oak in those regions during the LGM [48]. The modern-day northern-most limits of *H. maia* were not attained by oaks until about 10-11 ka [49], and *H. maia* likely arrived much later still, as the pollen of northern versus southern oak species cannot be separated. *Hemileuca maia* may have experienced a northward range shift during the Holocene Climatic Optimum between 5-8 ka [35], paralleling a northward shift of *Quercus*, with subsequent isolation of populations to microclimatically favourable sites such as sand and rock barrens as the climate cooled.

### Knowledge gaps and future research

Understanding pre-zygotic isolating mechanisms in the form of sex pheromone cross-compatibility offers a key insight to understanding *Hemileuca* evolution, guided by molecular variation and biogeography. Further research should be directed to establishing reproductive compatibility between *H. latifascia* versus *H. peigleri* and *H. slosseri*. Surprisingly, cross-trials of *latifascia* compared to Great Lakes poulations have never been examined, perhaps under the previous assumption that northern Great Plains populations represented *H. nevadensis*. Genetic analysis of Great Plains *H. latifascia* populations might be particularly informative, as the high diversity hinted at so far may shed light on patterns of glacial refugia. The genetic variation of the disjunct New Jersey *H. latifascia* population also remains unstudied, but the advance of genomic techniques coupled with more complete geographic sampling of the *Hemileuca maia* complex offers hope in resolving the taxonomy and evolutionary history of this complex and fascinating genus.

## Acknowledgements

I thank Jason Dombroskie, Sam Jaffe, Max Larrivee, Neil Schoppman, Cory Sheffield, Jim Troubridge, Nancy Wilson, and David Wagner for contributing to this work through providing additional information, voucher specimens and informative discussion. Jocelyn Gill and Christi Jaeger kindly provided technical support.

## Supporting information

**S1. *Hemileuca* voucher specimen data.**

